# Inhibitory mechanisms in the prefrontal-cortex differentially mediate Putamen activity during valence-based learning

**DOI:** 10.1101/2024.07.29.605168

**Authors:** Tal Finkelman, Edna Furman-Haran, Kristoffer C. Aberg, Rony Paz, Assaf Tal

**Author notes:** Equal contribution (Rony Paz, Assaf Tal).

## Abstract

Learning from appetitive and aversive stimuli involves interactions between the prefrontal cortex and subcortical structures. Preclinical and theoretical studies indicate that inhibition is essential in regulating the relevant neural circuitry. Here, we demonstrate that GABA, the main inhibitory neurotransmitter in the central nervous system, differentially affects how the dACC interacts with subcortical structures during appetitive and aversive learning in humans. Participants engaged in tasks involving appetitive and aversive learning, while using functional magnetic resonance spectroscopy (MRS) at 7T to track GABA concentrations in the dACC, alongside whole-brain fMRI scans to assess BOLD activation. During appetitive learning, dACC GABA concentrations were negatively correlated with learning performance and BOLD activity measured from the dACC and the Putamen. These correlations were absent during aversive learning, where dACC GABA concentrations negatively correlated with the connectivity between the dACC and the Putamen. Our results show that inhibition in the dACC mediates appetitive and aversive learning in humans through distinct mechanisms.

## Introduction

Learning from appetitive (gain) and aversive (loss) outcomes involves continuously monitoring one’s knowledge about the environment and employing behavioral adjustments whenever the outcomes differ from expectations. It is an essential process for behavioral adaptation to environmental changes (1). In humans, the neural representations of both appetitive and aversive stimuli converge on the dorsal ACC (dACC) (2–9) but with distinct neuronal patterns, potentially involving separate neural pathways or coding systems (10–13). The existence of different systems for encoding appetitive and aversive stimuli is further supported by single-neuron recordings from non-human primates, showing that distinct neuronal populations within the ACC respond separately to appetitive and aversive values (14–16). However, the differential neural mechanisms underlying these distinct populations remain poorly understood.

Animal studies suggest that inhibition influences learning by regulating synaptic plasticity, network dynamics, and the timing of neuronal activity in various brain regions, including the prefrontal cortex (17). In rodents, the connectivity between brain regions that respond to both gain and loss is regulated by the primary inhibitory neurotransmitter gamma-aminobutyric acid (GABA) (18). Another study showed that pharmacologically elevating the levels of GABA in the dACC impairs reward-based learning (19). Evidence in human studies indicates that GABA in the dACC is involved in a learning-based decision-making model (20,21). Specifically, it was reported that the basal concentration of GABA in the dACC negatively correlates with the integration of acquired information and with the brain activation measures from the dACC during reward-related decision-making tasks (22).

Based on this literature, we hypothesized that inhibition, particularly GABA, would differentially modulate brain activity during appetitive and aversive learning in human participants, and that interactions between inhibition, brain activity, and behavior (learning performance) should be evident in regions that traditionally support learning, such as the dACC, Striatum, Amygdala, and Insula (13,23–26). Direct evidence in humans would require measures of both GABA and brain activity during learning. Here, we used magnetic resonance spectroscopy (MRS), which allows the non-invasive measurement of GABA (27,28), and functional magnetic resonance imaging (fMRI) to measure brain activity. 117 participants performed probabilistic learning tasks under gain and loss conditions during MRS and fMRI scanning at a high (7T) fields strength. The MRS data were collected from the dACC using an optimized sequence that reliably separates the concentration of GABA and Glutamate (29), the main excitatory neurotransmitter, which can serve as a marker for cellular and neuronal activity (30,31).

## Results

### Game paradigm and learning behavior

MRS and fMRI were measured during four learning tasks (shown in Fig. 1A). In each task, participants had to select one out of two Chinese letters, after which corresponding feedbacks were presented. The four tasks differed in their win probabilities and their outcome types. In tasks with a 65-35% win probability (GP=65 game), the ‘best’ option provided winning with a 65% probability, while the other ‘worst’ option provided winning with a 35% probability. The GP=65 games are learning games because participants can learn there is a better option. As a control, we added an unlearnable condition, GP=50 games, with a winning probability of 50% for both options. Under the appetitive condition, winning was to gain money; under the aversive condition, winning was not losing money. For more details, see the materials and method section. Each game contained 50 trials, along which participants learned to choose to maximize their gains and minimize their losses.

**Figure 1.**
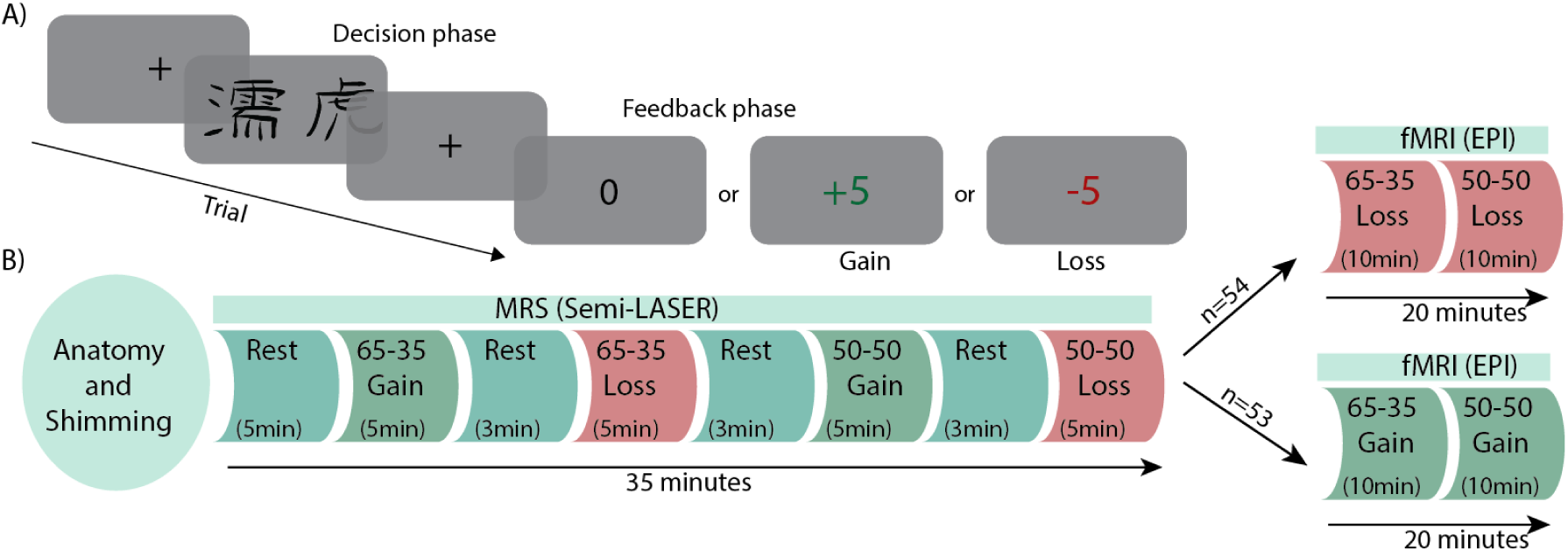
**A)** Behavioral paradigm and scanning protocol. Each trial starts with the presented frames-a waiting screen with a cross in the middle, followed by a two-letter frame where the participant has to choose one letter, followed by an another waiting frame (inter-stimulus-interval; ISI), and then the participant is presented with the outcome of their choice: 0 or +5 in the gain condition game and 0 or -5 in the loss condition game. **B)** The protocol inside the scanner. The MRS and MRI games were randomized. Half of the participants started with MRI blocks, and half started with MRS blocks. The numbers represent the probability for outcome.

The MRS portion of the task was comprised of eight blocks (Fig. 1B) that were collected in a sequence of four game-scans and four rest scans to allow GABA and Glu to return to baseline. The fMRI portion consisted of three blocks (one rest, two games) where half of the participants (n=53; 26 females) played two Gain games (GP=65-Gain and GP=50-Gain; Gain group), and the other half (n=52; 26 females) played two Loss games (GP=65-Loss and GP=50-Loss; Loss group). For the learnable GP=65 games, a learning score was defined as the probability of choosing the correct letter in the last 10 trials (i.e., the letter with a 65%-win probability). For the unlearnable GP=50 games, one letter was defined as the reference choice to which the learning score was referring.

Participants demonstrated learning and superior performance in the learnable conditions only (Fig.2A; PG=65; average learning scores: 0.709±0.002/0.701±0.002 in gain/loss), compared to the unlearnable conditions (Fig.2A; PG=50; 0.498±0.002/0.506±0.002 in gain/loss; ANOVA f{1,106}= 46.8; p-value = 5e-10; Table S1.). This shows successful and similar learning in both gain and loss conditions.

**Figure 2.**
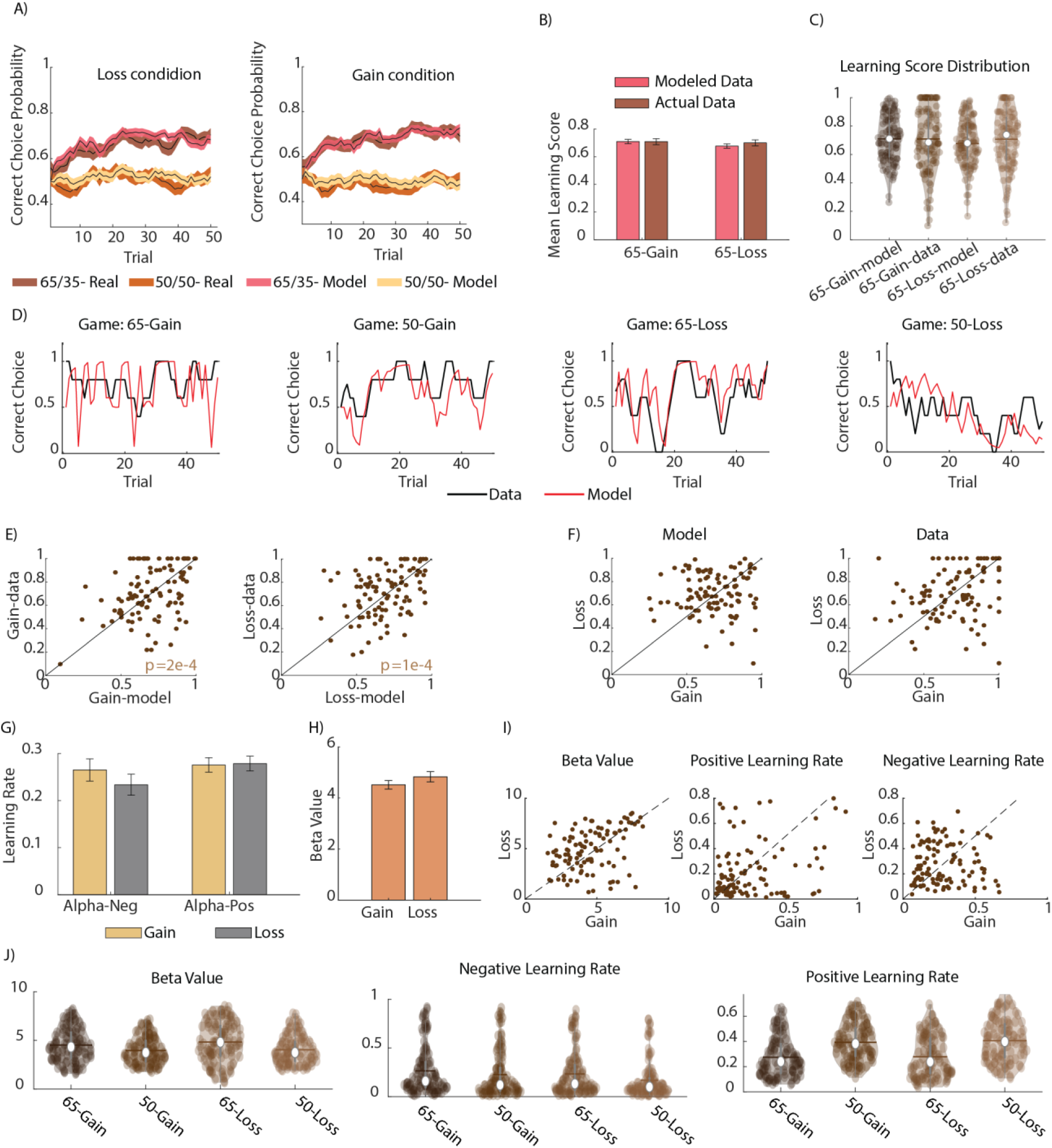
Behavioral data and model fitting. **A)** Learning curves for all four conditions. The shaded areas are SEM. **B)** Mean learning scores comparison. Error stands for SEM. **C)** mean learning score distribution. **D)** Single subject model fitting example. **E-I)** Model parameters are shown for learning conditions (65-gain and 65-loss). **E)** Comparison of the calculated learning scores in the original data and in the model. **F)** Learning scores comparison between gain and loss conditions. **G)** Mean Negative and positive learning rates (Alpha-Neg; Alpha-Pos). Error stands for SEM. **H)** Mean decision weight for expected value (Beta/β*_Q_*). Error stands for SEM. **I)** Model parameters comparison between gain and loss. **J)** Model parameters distribution for all game conditions.

We fitted a classical reinforcement-learning model (32) to the individual behavior for the purpose of obtaining the individual, per-trial, value difference coding for our fMRI analysis. The model contains a decision weight (β) parameter and separate parameters for positive (α+) and negative (α-) learning rates (See methods for more information). The learning scores were similar across model-fit and actual behavior for both gain and loss games (Fig. 2A-C; Model learning scores: Gain: 0.710±0.001; Loss: 0.677±0.001; Actual learning scores: Gain: 0.709±0.002; Loss: 0.701±0.002). Similarly, at the individual level, the learning scores of actual data and the model were strongly correlated in both gain and loss (Fig. 2E; Pearson correlation p-values: of 7 · 10^−9^ and 1 · 10^−8^, respectively), confirming that the model captures the variability in individual behavior (Fig. 2D depicts single-subject examples in each game condition). We did not find a significant difference in learning scores between gain and loss in either actual or modeled data (Fig. 2F, two-tailed t-tests). There were no interactions between model parameters (beta value and learning rates) and no interaction and changes across gain and loss conditions (Fig. 2G-J; repeated measures ANOVA test was conducted for each parameter with GP and LG as factors with no significant effects).

### MRS Data

We calculated the quality assurance parameters of the MRS signal (Fig. 3). The mean GM, WM, and CSF concentrations across subjects were 0.63±0.05, 0.28±0.06, and 0.08±0.03, respectively. The mean lipid/NAA ratio was 0.08±0.01, which indicates there are no extraneous lipids contaminating the spectra. The mean water FWHM and SNR are 14Hz ± 1Hz and 139±4). Taken together, these metrics indicate the spectral quality of the data.

**Figure 3.**
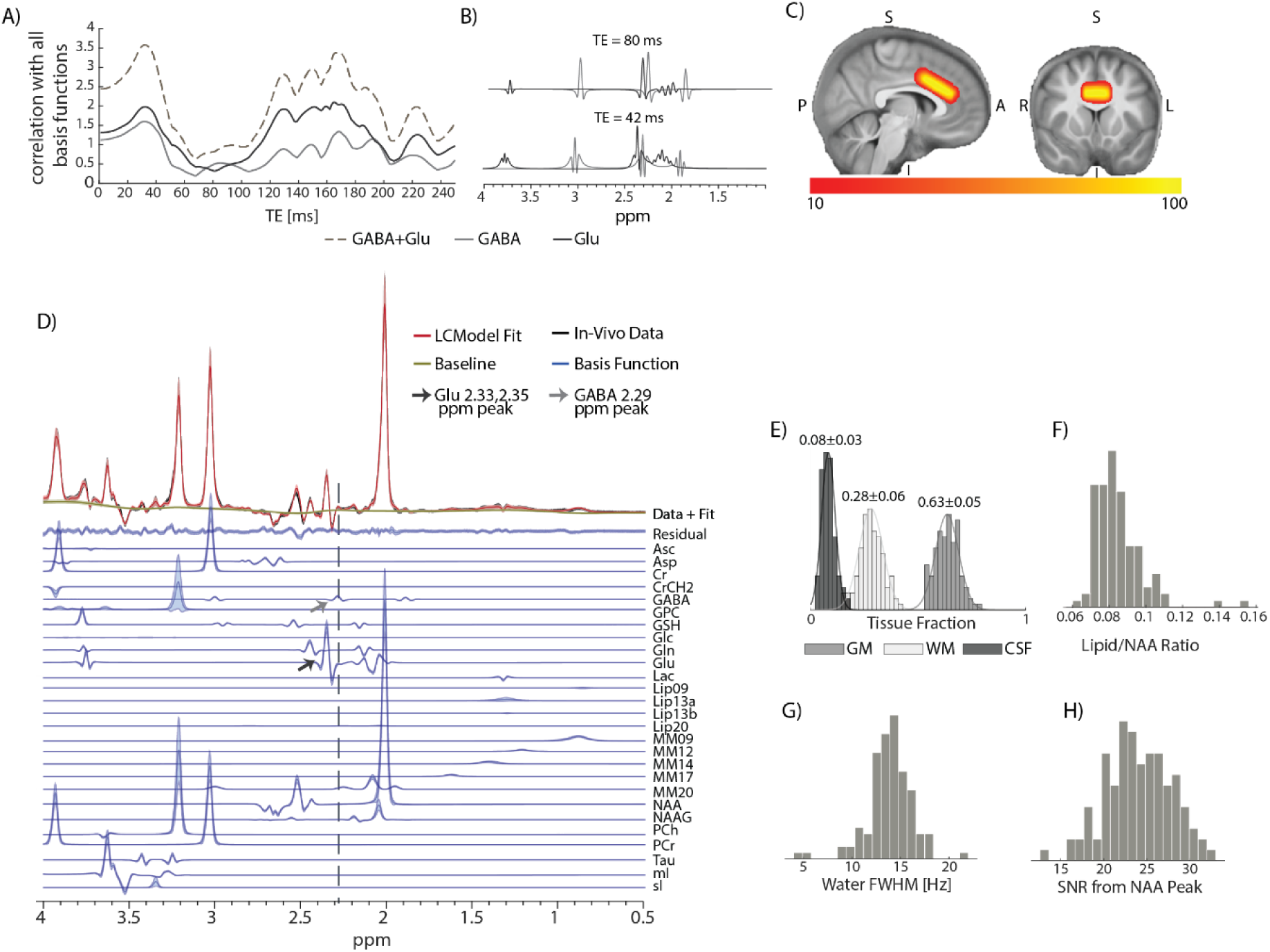
MRS spectral quality metrics. **A)** Simulation of basis function correlation with Glu, GABA, and Glu+GABA across different TEs (from TE= 1 ms to TE = 250 ms). The simulated metabolites are Asp, Asc, GPC, Cho, PCh, Cr, PCr, GABA, Glc, Gln, Glu, mI, Lac, NAA, NAAG, SI, GSH, and Tau. **B)** Glu and GABA basis function at TE=42ms and TE=80ms. Note the separable appearance at TE=80ms. **C)** Heat map of voxel positioning across all subjects. Voxels that are shared for less than 10 participants are filtered out. **D)** Mean LCModel Fitting across all subjects. **E)** Gray matter (GM), white matter (WM) and cerebrospinal fluid (CSF) distribution across all subject. mean±SD. **F)** Lipid/NAA ratio histogram across all subjects. **G)** Histogram of water FWHM at rest across all subjects. **H)** The histogram of SNR at rest was calculated using

### GABA differs between gain and loss conditions

The absolute concentrations of GABA, Glu, and the E/I balance (Glu/GABA) across conditions are presented in Table 1. We normalized the concentration during the games by subtracting the initial rest for inter-subject correlations. The normalized concentrations are denoted as ΔGABA and ΔGlu and ΔE/I balance (Table 2; see Table S2 for other metabolites). To examine the difference in the behavior of GABA in the gain and loss condition, we used a mixed model analysis (MM1) to model ΔGABA with the game probability (GP) and the game valence (gain or loss; GL). Overall, there was an interaction between GP and GL (GPxGL interaction: ANOVA f{1,291.3}= 4.8; p-value= 0.03, see MM1 in methods; Table S3, Fig. S1). During the loss condition, ΔGABA concentration in the learnable 65-Loss scenario was elevated compared to the 65-Gain game (Fig. 4A; two-tailed-t-test p=0.02), and marginally elevated compared to the 50-Loss (two-tailed-t-test p=0.08). During the gain condition, ΔGABA concentration did not change from the initial rest and was elevated during the unlearnable 50-gain scenario compared to the 65-gain condition (one-tailed t-test p=0.05). This is in accordance with our previous finding (33).

**Table 1.** Mean absolute concentration for GABA, Glu, and the E/I balance. The CRLB value refers to the accuracy of spectral fitting for the spectra acquired during the games. All errors presented are standard errors of the mean.

| Condition | GABA |  | Glu |  | E/I balance |
| --- | --- | --- | --- | --- | --- |
|  | Concentration [Mm] | CRLB (%) | Concentration [Mm] | CRLB (%) | Ratio Value |
| Initial Rest | 1.0143±0.02 | 17.57±0.3 | 12.667±0.1 | 3.20±0.04 | 12.49±0.1 |
| 65-Gain | 1.0075±0.02 | 17.79±0.3 | 12.798±0.1 | 3.23±0.05 | 12.70±0.1 |
| 50-Gain | 1.0479±0.02 | 17.70±0.3 | 12.881±0.1 | 3.22±0.05 | 12.29±0.1 |
| 65-Loss | 1.0626±0.02 | 17.56±0.3 | 12.814±0.1 | 3.26±0.04 | 12.06±0.1 |
| 50-Loss | 1.0188±0.02 | 17.79±0.2 | 12.824±0.1 | 3.23±0.04 | 12.59±0.1 |

**Table 2.**
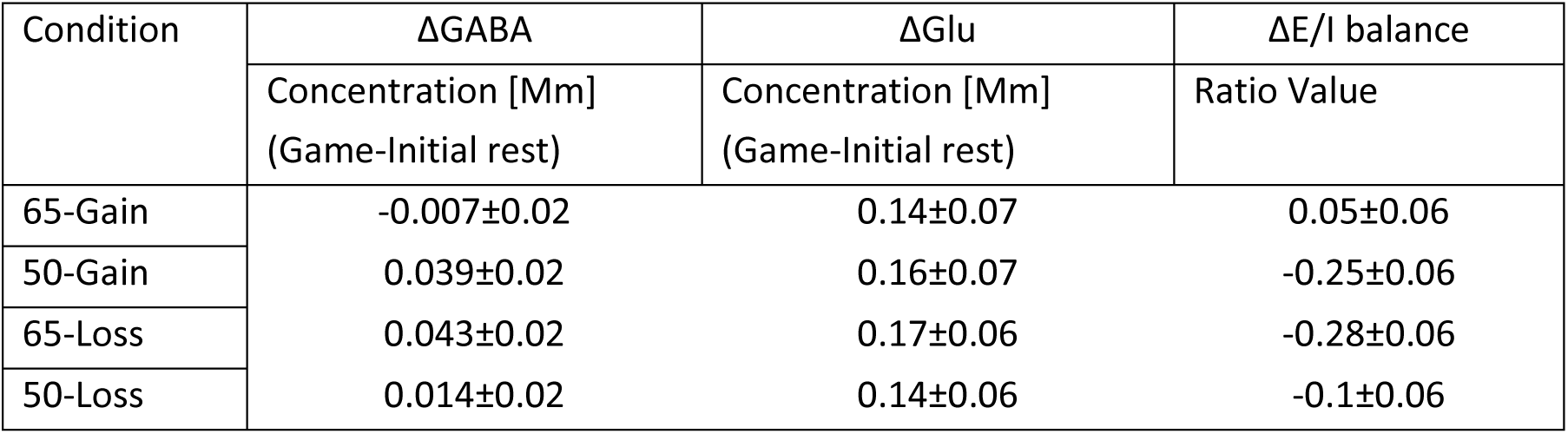
Mean concentration changes from the initial rest for GABA, Glu, and the E/I balance. All errors presented are standard errors of the mean.

**Figure 4.**
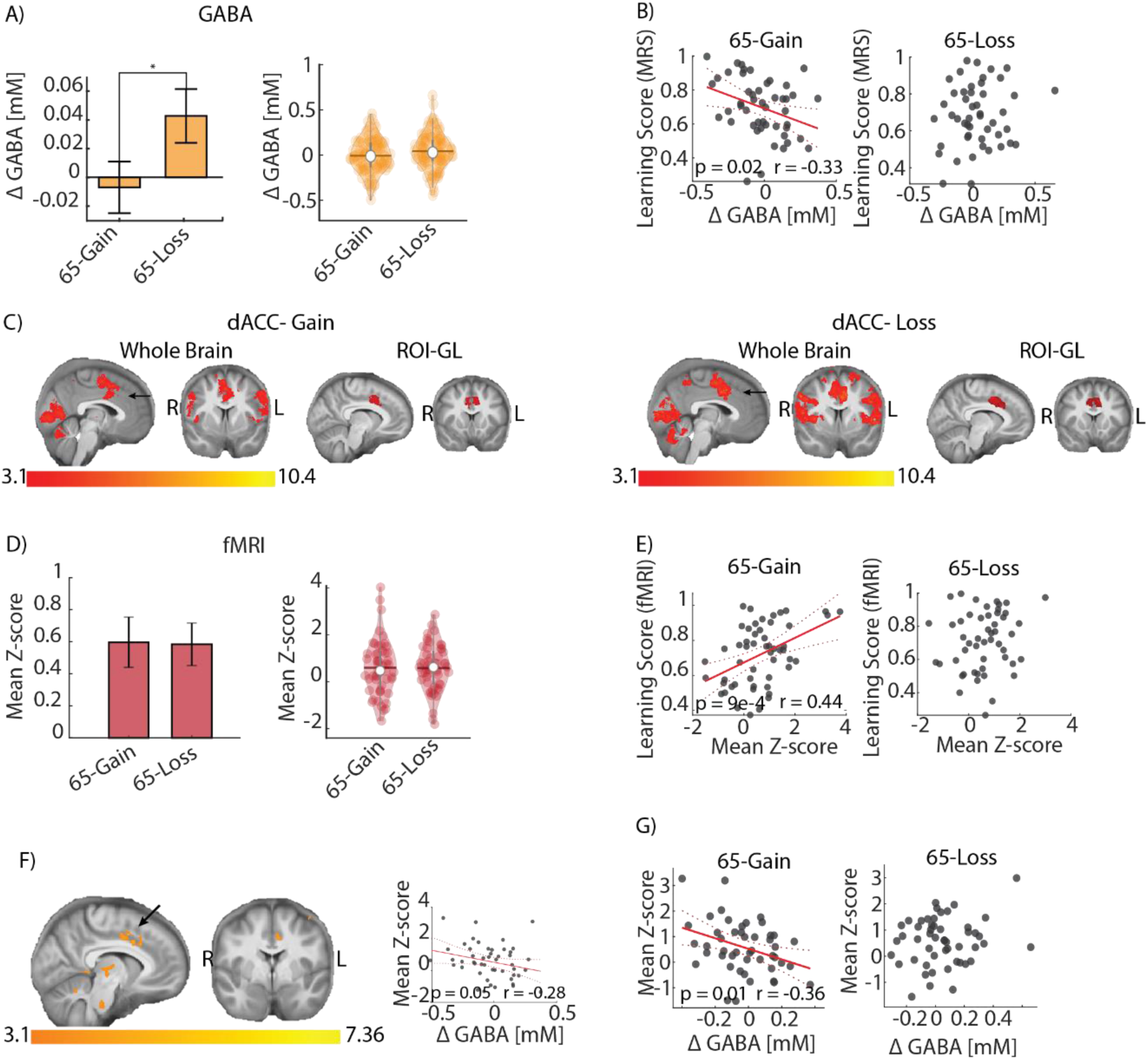
Correlation between ΔGABA, BOLD, and learning scores. **A)** ΔGABA mean with SEM and distribution in gain and loss. **B)** Correlation between the learning scores and ΔGABA from the dACC in gain and loss groups. **C)** Group-level activation of the dACC (GL map) and Group-level activation of the dACC within the spectroscopic voxel (ROI-GL map). **D)** BOLD mean Z score with SEM and distribution in gain and loss. **E)** Correlation between the learning score and the BOLD mean Z score from dACC in gain and loss groups. **F)** Right panel: Group-level activation of the learning correlates with BOLD in gain>loss (Learning-GL map). Left panel: Correlation between BOLD mean Z score from dACC-activated voxels (Learning-GL map) and ΔGABA from the dACC in gain. **G)** Correlation between BOLD mean Z score from dACC and ΔGABA from the dACC in gain and loss groups. ΔGABA represents GABA change from the initial rest.

We note that while ΔGlu consistently displayed an increase during all game scenarios and was correlated with ΔGABA (ANOVA f{1,380.4} =47; p-value= 3e-11; Table S3), there were no significant interactions or differences across game conditions for ΔGlu for ΔE/I balance (ANOVA for ΔGlu or ΔE/I with GP and LG as factors; p>0.1 for all). Therefore, we focus on ΔGABA interactions with behavior in this work.

### GABA correlates with learning under the gain condition

To explore the interplay between GABA concentrations and learning, we considered the Gain and Loss conditions separately (Gain group and Loss group). We tested the relationship between learning performance and ΔGABA (MM2 in methods; Table S4-S5) and found an effect in ΔGABA concentrations only within the gain group (Table S4; f{1,47}=4.7; p=0.02). Explicitly, at an individual level, ΔGABA concentrations were negatively correlated with learning performance (Fig. 4B; Gain: p=0.02; Pearson’s r=-0.33; Loss: p=0.8).

### BOLD in the dACC correlates with expected value and individual learning performance

We calculated the BOLD activity in the dACC using the spectroscopic voxel as a region of interest (ROI; ROI-GL map; Supp. Fig. S2). The design matrix used two explanatory variables (EVs): one for the decision phase, weighted by ΔQ (representing the expected value difference between selected and rejected options), and another for the feedback phase, weighted by PE (prediction error). These variables reflect neuronal coding of subjective value difference and prediction error, respectively (see material and method section for more details). Z-scored activation maps were generated for each EV, representing the activation probability of each voxel. In this work, we discuss only the decision phase contrast. To verify that the dACC activation is not dependent on our ROI analysis, we conducted a whole-brain analysis using the same parameters and found significant activation in the dACC (GL map; Supp. Fig. S3; Table S6).

BOLD activity in the dACC spectroscopic voxel during the decision phase did not differ across learnable game conditions (Fig. 4C, D, t-test p = 0.95; Supp. Fig. S1; Gain-65: 0.60+0.16; Gain-50: 0.68+0.15; Loss-65: 0.58+0.13; Loss-50: 0.49+0.14; ANOVA test, no effects). However, only under the learnable gain condition (PG-65 gain) the BOLD activity was positively correlated with individual learning scores (Fig. 4E; Gain: p = 9e-4, r= 0.44; Loss: p = 0.3, r= 0.1; cross-correlation coefficient difference (CCD): p= 0.05). See MM 3 in methods for more details (Table S7-S8; ANOVA f{1,47}= 9.1; p-value: 0.004).

We ran a third, separate group-level analysis (whole-brain) to verify this finding, using the individuals’ learning scores as a covariate. We found the dACC was significantly active in a gain>loss contrast (Learning-GL map; Table S9; Fig. 4F, left panel). This confirms that the dACC BOLD is correlated with learning during the gain but not the loss condition. Further, the correlation of this area with ΔGABA showed a negative correlation, similar to our finding using the ROI analysis (Fig. 4F, right panel; p = 0.05, r= -0.28).

### Activity in the dACC and Putamen is correlated with GABA during gain learning

Next, we evaluate the connection between the MRS and fMRI measurements. We applied linear regression (MM4 in methods; Table S10-S11) separately to the gain and loss group. We found a significant effect of ΔGABA only within the gain group (ANOVA f{1,77.5}= 5.1; p-value: 0.03, Supp. Fig. S1), showing a negative correlation between ΔGABA concentrations and BOLD activity within the dACC (ROI-GL). This negative correlation was specific to the learnable gain-65 condition (Fig. 3G, left panel: p=0.01, r=-0.36) and was not detected for loss (Fig. 3G, right panel; CCD: p= 0.003), or during the unlearnable conditions (Supp. Fig. S1). This provides a strong validation for our approach to measuring GABA and BOLD within subjects.

Subsequently, we examined the interactions between ΔGABA concentrations and the BOLD activity in other brain regions that contribute to valence encoding and decision-making processes (Table 3). We used the Harvard-Oxford subcortical structure atlas-based masks (M2-7). We found that BOLD activity in the left Putamen was negatively correlated with ΔGABA concentrations in the dACC only in the 65-Gain condition (Table 3. M5; p=0.02, r=-0.33; FDR corrected). To verify this finding, we tested it with a group-level map-based mask (GL map) and found the same result (Table 3. M9 Fig. 5A-B; p=0.02, r=-0.33). Additionally, the activity in the left Putamen was positively correlated with individual learning scores and only under the 65-Gain condition (Fig. 5A-B; p = 0.02, r= 0.33).

**Table 3.** All masks from which mean Z-score were extracted. dlPFC-dorsolateral prefrontal cortex; OFC-Orbitofrontal cortex; HOSSA-Harvard-Oxford subcortical structure atlas.

| Index | Mask location | threshold | Left and right<br>(separately) | Origin |
| --- | --- | --- | --- | --- |
| M1 | MRS functional voxel (dACC) |  |  | Cluster from ROI GL map |
| M2 | Insula | 5 | ✓ | HOSSA |
| M3 | Caudate | 5 | ✓ | HOSSA |
| M4 | Nucleus accumbens | 5 | ✓ | HOSSA |
| M5 | Putamen | 5 | ✓ | HOSSA |
| M6 | Amygdala | 5 | ✓ | HOSSA |
| M7 | OFC | 5 | ✓ | HOSSA |
| M8 | dIPFC |  | ✓ | Cluster from GL map |
| M9 | Left Putamen |  |  | Cluster from GL map |
| M10 | Left Insula |  |  | Cluster from GL map |

**Figure 5.**
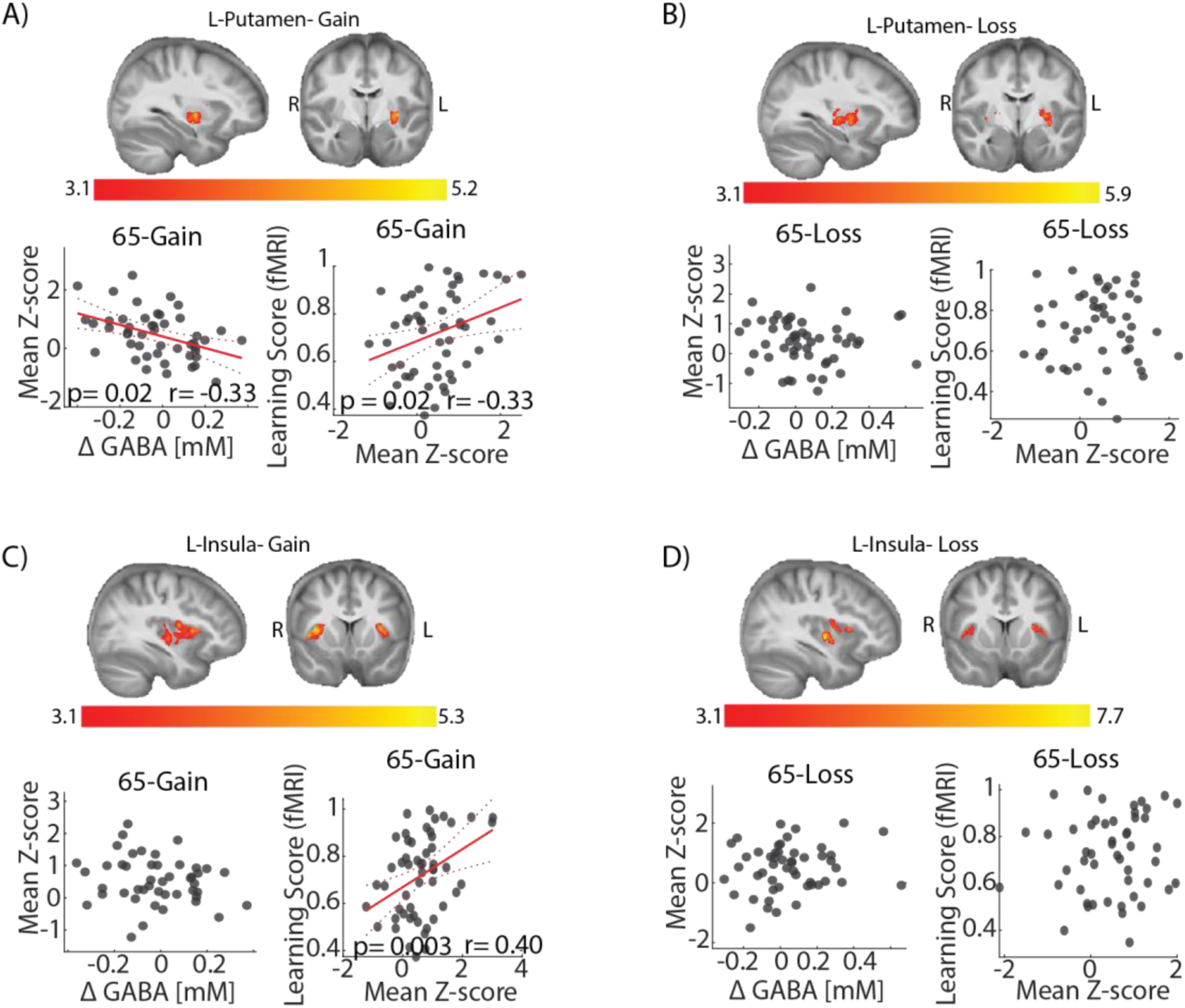
Learning and ΔGABA correlation in the Putamen and the Insula. **A-B)** Correlation between BOLD mean Z score from the left Putamen cluster of the group analysis and ΔGABA or learning score in gain (A) and loss (B) groups. **C-D)** Correlation between BOLD mean Z score from the left insula cluster of the group analysis and ΔGABA or learning score in gain (C) and loss (D) groups.

Moreover, we inspected on the correlation with the Insula, which forms the salience network together with dACC (34). We did not find a correlation between ΔGABA in the dACC and the BOLD in either the left or right Insula. This was examined on an atlas-based mask (Table 3. M2) and a group-level map (Table 3. M10) for presentation purposes (Fig. 5C-D). However, we found a significant positive correlation with the learning score under the 65-Gain condition only (p = 0.003, r= = 0.40), supporting that the salience network participates in decision-making guided learning.

### dACC-Putamen connectivity is correlated with GABA during loss learning

Beyond direct modulation of activity, inhibitory mechanisms can also modulate information transfer between the dACC and other brain regions. To test this hypothesis, we examined the relationship between ΔGABA in the dACC and BOLD-based functional connectivity. First, we calculated the connectivity during gain and loss between the dACC and several decision-making-related brain areas (Fig. 6A), as well as other reference areas (Table S12). We found significant functional connectivity between the dACC and the Putamen, as well as the dlPFC, nucleus accumbens, thalamus, left amygdala, and insula (Bonferroni correction for multiple comparisons). As in previous studies, there was strong connectivity between the dACC and the right and left insula (35).

**Figure 6.**
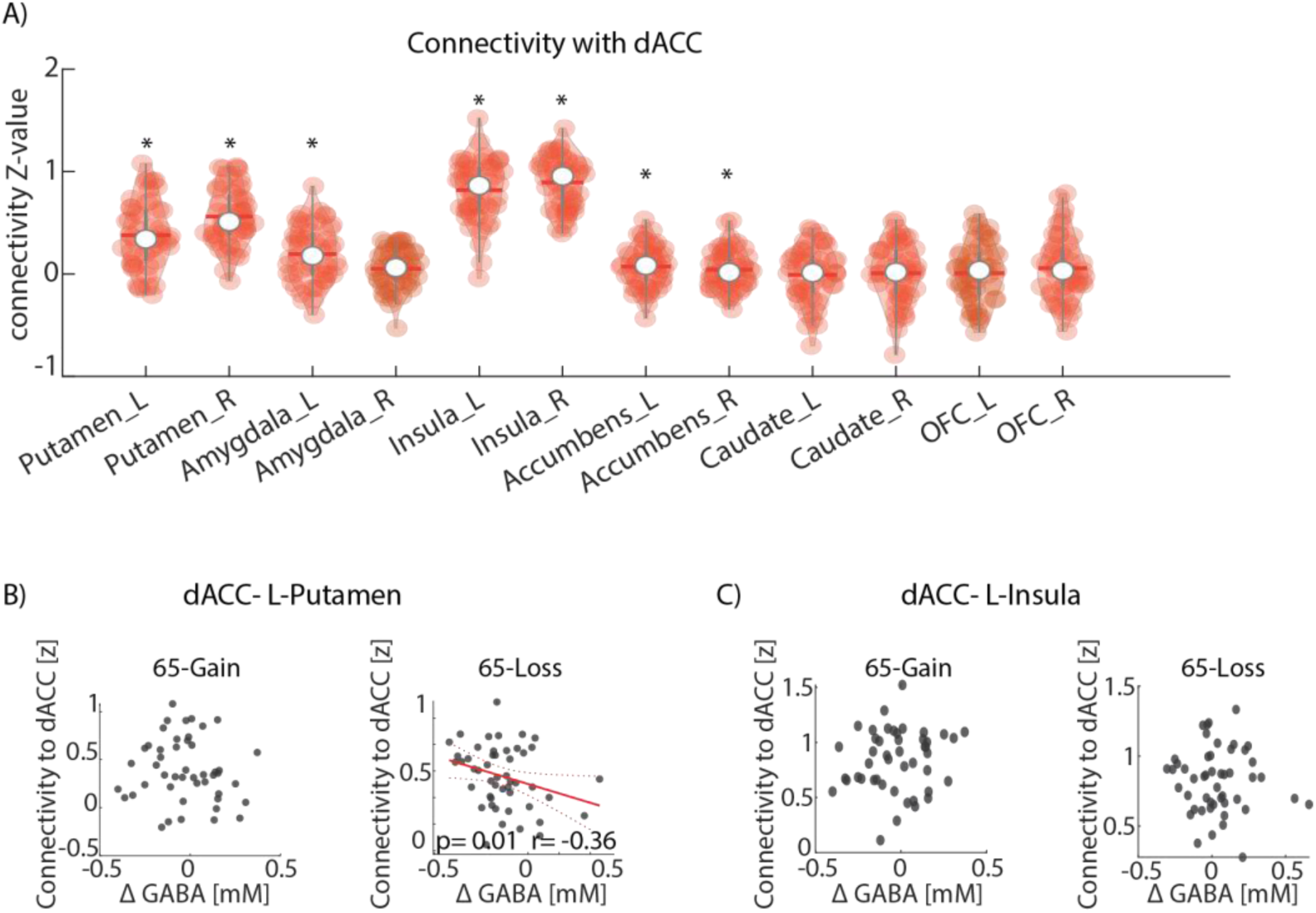
Correlation between connectivity and ΔGABA. **A)** Connectivity strength distribution for the connectivity between the dACC cluster of the ROI group analysis and the listed regions under the gain condition. **B)** Correlation between the left-Putamen-dACC connectivity in z value and dACC’s ΔGABA in gain and loss. **C)** Correlation between the left-insula-dACC connectivity in z value and dACC’s ΔGABA in gain and loss.

However, and importantly, ΔGABA concentrations in the dACC were negatively correlated with the strength of connectivity between the dACC and the left Putamen only and during the loss-65 condition only (Fig. 6B, p = 0.01, r = -0.36; FDR correction to all brain regions listed in Table S12). This implies that dACC GABA reduces the dACC-Putamen connectivity during loss.

## Discussion

Several studies have proposed that the ACC encodes positive and negative valences and salience within distinct sub-regions (15,23,36–38). This has been demonstrated in primates, which show an anatomical intermixing of gain and loss-related neurons in the ACC, with only a minority of neurons showing excitatory signaling for both gain and loss (14,16,39). Similarly, evidence from BOLD-fMRI studies point to distinct inhibitory neuronal mechanisms for appetitive and aversive learning in the dACC (11). The current study provides direct evidence that GABA concentrations in the dACC differentially mediate aversive- and appetitive-based learning in behaving humans. Under the appetitive condition, GABA concentration is negatively correlated with learning performance, and is associated with the BOLD signal that encodes values in both the dACC and the left Putamen. Additionally, the BOLD signal in the dACC and the left Putamen shows a positive correlation with learning performance in both regions. Hence, the greater the inhibition, the weaker the activity in the dACC and the left Putamen, and the lower the learning score. However, under the aversive condition, GABA is uncorrelated to behavioral or physiological measures, except for a negative correlation with the dACC-Putamen connectivity. These two distinctive patterns in appetitive and aversive learning conditions are consistent with the presence of two neuronal populations: one that controls appetitive value coding via projection from the dACC to the left Putamen and another related to aversive value coding via a different yet-undetermined mechanism. This is also consistent with the size of our spectroscopic voxel (10 mL), which is large enough to contain such a heterogeneous population of neurons. Moreover, GABA concentrations during the loss condition are elevated compared to the gain condition, which might point toward a mechanism of inhibitory saturation in the loss condition that hinders the gain-related neurons (40,41).

The observed negative correlation between GABA concentration changes from the baseline (ΔGABA) and reward-guided learning scores is consistent with previous MRS literature, which has largely demonstrated negative correlations between task-related GABA concentration changes and various learning-related performance scores (42–45). One possible explanation for this negative correlation could be tied to the association of increased GABA concentrations in the ACC with exploration under reward-related tasks (19,21,46). In our paradigm, the best strategy would be to find the better letter by exploring both options and then choosing it repeatedly, so increased exploratory behavior might not have been advantageous within our paradigm; it could explain the negative correlation we observed. A study using a similar paradigm to ours and measured rest concentrations of dACC GABA showed a positive correlation between GABA concentration at rest and reward-guided task performance (47). We find a comparable correlation between basal GABA concentration and learning performance (Supp. Fig. S4). The explanation for the differences in the correlations between GABA concentration at rest and ΔGABA concentration lies in our definition of ΔGABA as the subtraction of GABA concentration at rest from GABA concentration during the game.

ΔGlutamate concentrations consistently increased across all game conditions (Table 2), indicating ongoing dACC activity throughout these scenarios, driven by both metabolic (from the citric acid cycle) and neurotransmitter dynamics (30,31). This is supported by elevated BOLD measurements and heightened lactate concentrations (Supplementary Table S2) across all game conditions, consistent with increased oxidative metabolism (48). These observations parallel earlier research demonstrating elevated Glu concentrations in the ACC during cognitive tasks and studies highlighting correlations between Glu dynamics, BOLD response, and task timing (48–52). The observed elevation of GABA might also explain elevated Glu, as Glu is a precursor for its synthesis (53); however, in this case, we would expect to see differences in ΔGlu between the gain and loss conditions, which we do not. Additionally, we did not find a relationship between ΔGlu and BOLD measures during learning, suggesting that the increase in ΔGlu is not necessarily related to the measured value coding-related BOLD changes. Currently, it is hard to conclude whether this finding merits interpretation or is due to lower MRS sensitivity and/or volume averaging over functional subnetworks within the dACC.

Previous findings and theoretical models suggest that the E/I balance plays a major role in brain activity, cognition, and behavior (54–57). We employed a simulated likelihood ratio test (SLRT) to compare our mixed models of ΔGABA with identical models featuring ΔE/I balance instead of ΔGABA (MM1-4) to examine if the effect we detected is related to GABA or to the E/I balance (Table S12). We found that the ΔE/I balance offers a poorer fit to the game conditions and learning scores (mixed models 1 and 2). This could be attributed to the metabolic contribution to ΔGlu (31), potentially influencing the actual ΔE/I balance value. Alternatively, our findings may indicate a distinct role of GABA, separated from the E/I balance context, during the value difference encoding phase, particularly in actions guided by learning.

We noticed a negative correlation between GABA concentrations and BOLD activity in the human left Putamen during appetitive – but not aversive – value difference encoding, implying that the increased inhibition in this condition suppresses a projection from the dACC to the left Putamen (Table 3: M2-M10). This conclusion is supported by primate studies, which show that the Putamen-caudate complex, receiving input from the ACC, plays a role in processing gain-related signals rather than loss-related ones (58,59). Additionally, mice studies show that increased inhibition during appetitive tasks might influence neuronal populations that rely on previous information for decision-making (19,46). The absence of this relation between the dACC and the putamen under the aversive condition is consistent with previous reports that aversive and appetitive learning is driven by different neuronal populations (14–16,60). However, our data is insufficient to explain the underlying differential mechanism nor pinpoint additional regions that drive it.

One possible explanation for the aversive learning-related increase in ΔGABA might originate from a neuronal population that impedes the interaction between the dACC and the Putamen. This is supported by the intriguing finding that the salience network exhibited the strongest connectivity during both gain and loss, aligning with existing research suggesting its involvement in gathering essential information for decision-making processes (61,62) (Table S11); As well as the negative correlation between ΔGABA concentrations in the dACC and the strength of connectivity between the dACC and the Putamen during the loss – but not gain – condition.

In this study, we compared MRS data acquired during separate gain and loss reinforcement-learning games. However, a paradigm that includes mixed motivation components, linking aversive and appetitive stimuli, is recommended to isolate aversive motivation mechanisms more effectively (63,64). This suggests that our paradigm may not be optimal for detecting such mechanisms and might explain why we did not observe differences in BOLD activity or the learning performance between the gain and loss conditions. Additionally, our MRS and fMRI data were acquired separately under the assumption that task-induced changes are similar in subsequent games. To overcome these limitations, we suggest implementing an event-related paradigm combining reward and punishment and using an interleaved MRS-fMRI sequence(65–69). Alterations to metabolite transverse relaxation times (T2) due to the crossing of neurotransmitters between intravesicular, intracellular, and extracellular pools might act as a confounding factor in our analyses. This could be mitigated in future studies by implementing multiparametric fMRS sequences (70–74), which would simultaneously quantify metabolite relaxation times and concentrations throughout the rest and all task conditions.

## Materials & Methods

### Subjects

We enlisted 117 healthy volunteers (56 females; mean Age: 27±5 years) without pre-existing psychiatric and neurological conditions. All participants provided written informed consent under the approval of the Wolfson Medical Center Helsinki Committee (Protocol number: 0084-19-WOMC). The protocol was further approved by the ethical review board of the Weizmann Institute of Science. Five participants were excluded due to excessive movement and one for non-completion of the task. Among the remaining 111 volunteers, five have missing MRS data, while four underwent only MRS acquisition. 105 participants have full data from both fMRI and MRS acquisitions.

### Experimental Design

A decision-making task paradigm (Fig. 1A) was used in this study in four conditions. During each task, in every trial, participants selected one out of two Chinese letters, after which corresponding feedback was presented. The four tasks differed in their winning probabilities and in their outcome types. Each game included 50 trials. In tasks with a 65-35% win probability (GP=65 game), the ‘best’ option provided winning feedback with a 65% probability, while the other ‘worst’ option provided winning/losing feedback with a 35% probability. The GP=65 games are called learning games as participants can learn there is a better option. As a control, an unlearnable condition was included, GP=50 game, where winning feedback was presented with 50% probabilities for both options. Under the appetitive condition, winning was gaining money; under the aversive condition, winning was not losing money. In summary, there were two Gain games (GP=65-Gain, GP=50-Gain) and two Loss games (GP=65-Loss, GP=50-Loss). Participants were told to maximize their scores. The game presentation was implemented in MATLAB (R2021b) using the Psychophysics Toolbox (75,76). All participants had a training session before entering the scanner.

The MRS scan was comprised of eight blocks (Fig. 1B) that were collected in a sequence of four game-scans and four rest-scans in the following order: baseline (rest, 5min), game 1 (5min), rest (3min), game 2 (5min), rest (3min), game 3 (5min), rest (3min), game 4 (5min). The type of game presented was randomized. The rest of the scans were interspersed between game scans to allow GABA and Glu to return to baseline.

The fMRI scan consisted of two blocks (two games-10 minutes each) where half of the participants (n=53; 26 females) played two Gain games (GP=65-Gain and GP=50-Gain), and the other half (n=52; 26 females) played two Loss games (GP=65-Loss and GP=50-Loss). The beginning of each game was synchronized to the beginning of the first TR, using a trigger command. The inter-stimulus interval (ISI) and inter-trial interval (ITI) are defined as a random exponential distribution around 4.5sec and 6sec with a minimum value of 2.5sec and 4sec for the fMRI games, and 2.5sec with a minimum value of 0.5sec and 1sec for the MRS games, to keep the game length at 5 minutes. For the learnable GP=65 games, a learning score was defined as the probability of choosing the correct letter in the last 10 trials (i.e., the letter with a 65%-win probability). For the unlearnable GP=50 games, one letter was defined as the reference choice to which the learning score was referring.

### Behavioral modeling

We used a common Q-learning model that was previously shown to be a good model for our kind of task (32). In this model, the expected value *Q_i_*(*t*) of the selected letter *i* in trial *t* was updated by the mismatch between the expected value and the actual outcome *R(t)*, i.e., the prediction error- δ*_Q_*(*t*), scaled by a learning rate α. Separate learning rates were defined for positive prediction error (α_*PE+*_) or negative prediction error (α_*PE−*_):

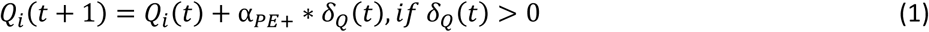

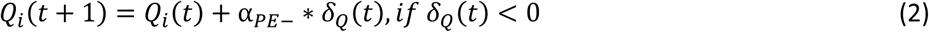

The probability *p* of selecting a particular letter *i* depends in trial *t* on its utility, u_*i*_ (SoftMax choice-probability):

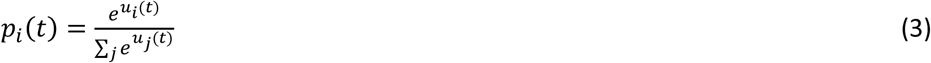

Here, u_*i*_ is defined as the sum of parameters that may contribute to a decision scaled by their respective decision weights. For example, the decision weight β*_Q_* determined to what extent decisions are decided by expected values, such that large and small values of β*_Q_* Indicated that decisions are more and less influenced by expected values, respectively:

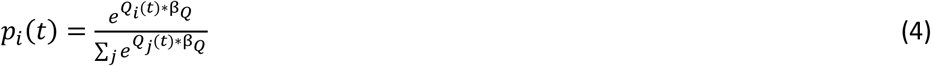

In order to compare between the games, we modeled each game separately for each subject.

### MR Protocol

The scans took place using a 7T Terra scanner (Siemens Healthineers, Erlangen, Germany) and a commercial single-channel transmit/32-channel receive head coil (NOVA Medical Inc., Wilmington, MA, USA), reaching a maximum B1+ amplitude of 25 µT.

A T1 weighted image acquired with MP2RAGE (Magnetization Prepared 2 Rapid Acquisition Gradient Echoes; TR/TE/TI1/TI2=4460/2.19/1000/3200 ms, α_1_=4°/α_2_=4°, 1 mm^3^ isotropic voxels, TA=6:56 min). An initial localizer and gradient echo field map were acquired for automated B_0_ shimming (Scan parameters for B_0_ mapping: TR/TE_1_/TE_2_=232/3.06/4.08 ms, α=25°, 40 2.5 mm axial slices, 2.0×2.0mm^2^ in-plane resolution, TA=0:31 min) before MP2RAGE.

A multiband T2*-weighted gradient echo-EPI sequence developed at Minnesota, was used for fMRI data collection with the following parameters: TE=22.22ms, TR=1000ms, flip angle=45°, slice thickness of 1.6mm, no interslice gap, in-plane resolution 1.6 × 1.6 mm^2^, 130-time points, dwell time of 0.32sec. A multi-band factor of 5 (slice direction), parallel imaging (GRAPPA) factor of 2 (phase direction), phase partial Fourier-7/8. Four dummy scans, followed by a higher resolution volume collection for registration improvement, were executed prior to the EPI acquisition (77,78).

MRS data were acquired using an optimized sLASER sequence (TR/TE=7000/80 ms), which can detect both GABA and Glu with high precision due to a minimal spectral correlation (Fig. 3A) and a pseudo singlet appearance (Fig. 3B) (29). A 25X40X10 mm^3^ voxel was manually placed on the dACC (Fig. 3C) using anatomical markers. The maximal amplitude is set to 25 mT/m, dwell time of 0.25 ms, spectral BW of 4000 Hz, and 1024 data points.

### MRS Analysis

VDI libraries were utilized for all preprocessing steps (79). This encompassed coil combination, alignment, and phase correction through a well-established iterative algorithm (80).

To segment the T1-weighted anatomical images into gray matter (GM), white matter (WM), and cerebrospinal fluid (CSF) images, SPM12 from the Welcome Center for Human Neuroimaging, UCL, UK (http://www.fil.ion.ucl.ac.uk/spm), was employed (81). VDI was also used to calculate tissue fractions within the spectroscopic voxel for subsequent signal quantification.

LCModel was used for metabolite quantification (Fig. 3D). The basis set contained 17 metabolites in their basis sets: aspartate (Asp), ascorbic acid (Asc), glycerophosphocholine (GPC), phosphocholine (PCh), creatine (Cr), phosphocreatine (PCr), GABA, glucose (Glc), glutamine (Gln), Glu, Myo-inositol (mI), lactate (Lac), N-acetyl aspartate (NAA), N-acetylaspartylglutamate (NAAG), scyllo-inositol (Scyllo), glutathione (GSH), and taurine (Tau). Absolute quantification was carried out by correcting the metabolite concentrations provided by LCModel for tissue fractions estimated from the segmented images (82) and for relaxation effects, using literature values of T_2_ (Supplementary Table S14). We assumed the same T_2_ for gray matter (GM) and white matter (WM) fractions and no metabolites in cerebrospinal fluids (CSF) tissue fractions (83). Metabolites for which T_2_ relaxation data was not found in the literature were not quantified or further considered. The long TR eliminated saturation effects. Therefore, no T_1_ corrections were required. For each metabolite, concentrations that were three median deviations away from the median were excluded.

### MRS spectra quality assurance metrics

To evaluate lipid contamination within the spectrum, we determined the ratio of lipid signal intensity spanning 0.3-1.7 ppm to the NAA signal intensity spanning 1.9-2.1 ppm. The signal-to-noise ratio (SNR) was computed by dividing the NAA peak signal within 1.9-2.1 ppm by the standard deviation (STD) of the noise spectrum. The noise STD was derived from the central half of the spectrum, excluding its first and last quartiles. Water FWHM was quantified as the width of the water peak at half of its maximum height in Hz, enabling assessment of spectral linewidth.

### fMRI Analysis

fMRI data were preprocessed using FEAT (FMRI Expert Analysis Tool) version 6.0.4, part of FSL (FMRIB’s Software Library, www.fmrib.ox.ac.uk/fsl). Preprocessing steps included motion correction, slice timing correction, temporal high-pass filtering (0.008 Hz), and pre-whitening (84). Head motions were corrected using MCFLIRT (85). Non-brain areas were removed using BET. Each image was then spatially smoothed using a Gaussian kernel of FWHM 4 mm. Next, functional and structural images were co-registered to a standard (MNI152 atlas) image space.

First-level statistical analysis was done using a general linear model (GLM). The design matrix included two weighted modeling explanatory variables (EVs) – the decision phase, weighted per trial by the ΔQ value (the expected value difference for the selected option minus the rejected option-indicating the neuronal coding of subjective value difference), and the feedback phase, weighted per trial by the PE value-indicating the neuronal coding of subjective PE), their time derivatives, and extended motion parameters EVs (86). We used a measure of frame-wise displacement to find and remove trials with excessive head movement as confound (i.e. EVs with a threshold of 0.9 mm). Trials with no response to the task were also removed from the analysis. For higher-level analysis, the single-subject functional maps were registered in the Montreal Neurological Institute (MNI) 152 space (87). Z-scored activation maps were generated for the decision phase and the feedback phase, representing the activation probability of each voxel. In this paper, we discuss only the decision phase contrast.

Three Group-level analyses are performed. First, we contrast BOLD responses for decisions made in Gain and Loss games separately, using an a priori ROI and a mixed effect FLAME 1+2 with a cluster-based threshold of Z=3.1 and p=0.05 (88). The ROI was defined as the mean MRS voxel (ROI-GL map; Supp. Fig. S2). Within the ROI mask, voxels shared by less than 5 subjects (out of 105) were excluded. Second, we test the same contrast for the whole brain (GL map; Supp. Fig.S3). Finally, we correlate differences in learning scores (i.e., Gain minus Loss) with the BOLD responses for Gain versus Loss games. The learning scores were demeaned and defined as a continuous covariate (Learning-GL map).

Mean Z-score, parameter estimates, and time series were extracted using Featquery (http://www.FMRIb.ox.ac.uk/fsl/feat5/featquery.html) from regions associated with different decision-making processes such as prediction error, expected value, and incorporation of learned information (for a complete list of regions, see Table 3) (11,89–91). The Harvard-Oxford probabilistic atlas in FSL was used to obtain anatomic masks of the caudate, insula, nucleus accumbens, orbitofrontal cortex, Putamen, amygdala, thalamus, subcallosal cortex, and posterior cingulate cortex (PCC). A mask of the dlPFC was derived from the whole brain group-level thresholded map, as we did not find it in the Harvard-Oxford atlas. Masks for the left insula and left Putamen were derived from the whole brain group-level thresholded map for presentation and finding verification purposes.

### Functional connectivity analysis

Beta-series correlations were used to measure the connectivity during the task (92). We conducted separate analyses for stimuli events and outcome events. We calculated the connectivity between the dACC cluster from the ROI group analysis and 24 other brain areas (Table S12); some are decision-making-associated, and others are not, for reference. We used partial correlation with the mean whole-brain beta series to account for global signal changes. The correlation coefficients were transformed into z-score and were used for the group-level connectivity estimation. We used Bonferroni correction for multiple comparison correction (N = 24).

### Mixed Linear Models

Mixed linear models were calculated to assess the correlations with the mean Z score, ΔGABA, and the learning score. Four models were calculated, and all of them included the subjects’ random effect. The first model explains ΔGABA with the game probability (GP) and reinforcers (gain or loss; GL) with an option of interaction and ΔGlu.

Mixed model 1 (MM1):

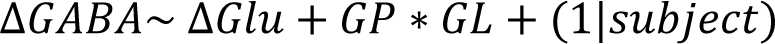

The second model explains the learning score during the MRS games with ΔGABA and ΔGlu during the learnable 65-35 game. Separate models were calculated separately for the gain group and for the loss group.

Mixed model 2 (MM2):

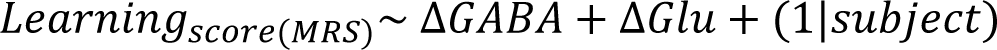

The third model explains the mean Z score during the decision phase with ΔGABA, ΔGlu, and the learning score during the 65-35 fMRI task. Separate models were calculated separately for the gain group and for the loss group, as we do not have all variables for each subject.

Mixed model 3 (MM3):

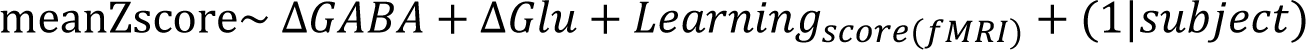

The fourth model explains the average Z score in the decision phase using only the changes in ΔGABA and ΔGlu, along with their interaction with game probability, in order to investigate an influence that could potentially be obscured by the correlation with learning. Distinct models were computed for both the gain and loss groups.

Mixed model 4 (MM4):

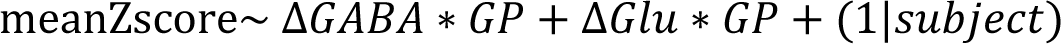

### fMRI correlation with MRS data

In order to evaluate the connection between MRS and fMRI measurements, we employed linear regression. This involved correlating the average brain activities, represented in Z-score values, or the mean connectivity coefficients derived from various brain regions with ΔGABA or ΔGlu (normalized concentrations), which served as the variables of interest in this analysis. These calculations were conducted independently for both the gain and loss groups.

Although MRS and fMRI data were acquired separately for each game condition, we consider these datasets comparable because they were acquired within a subject and within the same session and because participants practiced the games prior to scanning. Indeed, there was no significant change in learning scores between games during MRS and fMRI acquisitions.

As part of our quality control process, participants were excluded using the following method: Each data point’s y-value was compared to a normal distribution centered around a specific x-value. If the likelihood of obtaining such a measurement was less than 1%, the data point was excluded from the analysis (p < 0.01).

## Acknowledgments

Dr. E. Furman-Haran is grateful for the Calin and Elaine Rovinescu Research Fellow Chair for Brain Research and extends acknowledgments to Dr. Sagit (Wolfson Medical Center) for the Human MRI studies. Dr. K.C. Aberg acknowledges the Sam and Frances Belzberg Research Fellow Chair in Memory and Learning. We extend our appreciation to the Center for Magnetic Resonance Research (CMRR), University of Minnesota, USA, for providing the pulse sequences. Special thanks go to Edward J. Auerbach, Ph.D., and Małgorzata Marjańska, Ph.D. (CMRR) for developing the spectroscopy pulse sequence.

## Author Contribution

Conceptualization: TF, AT, RP

Methodology: TF, AT, RP, KCA, EFH

Investigation: TF, KCA

Visualization: TF

Supervision: AT, RP

Writing—original draft: TF

Writing—review & editing: TF, AT, RP, KCA, EFH

## Data and material Availability

All data supporting the findings of this study are available from the corresponding author upon reasonable request. Raw data are available from the corresponding author upon reasonable request. Custom code, FSL results maps, and processed parameters for behavioral, MRS, and fMRI tests are available at Dryad: https://datadryad.org/stash/share/lkmGBjijMzOX71mxz38FgPDFOX479bZfTCm7wmedunw.

