## Supplemental tables and figures for "Inhibitory mechanisms in the prefrontal-cortex differentially mediate Putamen activity during valence-based learning"

[4]Department of Biomedical Engineering, Tel Aviv University, Tel Aviv, Israel

† Equal contribution

### Supplementary data

**Table S1. fMRI learning score model results.** Repeated measures ANOVA on the modeled learning score in the fMRI tasks with LG and GP as factors. LG- boolean loss/gain variable. GP- game probability, Boolean: 50% is 1, and 65% is 0.

| Variable | f value {Df1, Df2} | pValue |
|---|---|---|
| Intercept | 862.1 {1, 209} | 3e-76 |
| LG | 0.02 {1, 209} | 0.9 |
| GP | 46.8 {1, 106} | 5e-10 |
| LG*GP | 0.07 {1, 106} | 0.8 |

**Table S2. Metabolites mean concentration changes from the initial rest.** The CRLB value refers to the accuracy of spectral fitting for the spectra acquired during the games. All errors presented are standard errors of the mean.

| Condition | tNAA | | tCr | | tCho | |
|---|---|---|---|---|---|---|
| | ΔConcentration [Mm] | CRLB (%) | ΔConcentration [Mm] | CRLB (%) | ΔConcentration [Mm] | CRLB (%) |
| 65-Gain | 0.024±0.009 | 1.000±0 | 0.06±0.03 | 1.000±0 | 0.10±0.04 | 1.01±0.03 |
| 50-Gain | 0.01±0.01 | 1.000±0 | 0.02±0.03 | 1.000±0 | 0.06±0.04 | 1.01±0.02 |
| 65-Loss | 0.01±0.01 | 1.000±0 | 0.04±0.03 | 1.000±0 | 0.10±0.04 | 1.01±0.02 |
| 50-Loss | 0.024±0.009 | 1.000±0 | 0.07±0.03 | 1.000±0 | 0.09±0.04 | 1.01±0.02 |

| Condition | mI | | Lac | | GSH | |
|---|---|---|---|---|---|---|
| | ΔConcentration [Mm] | CRLB (%) | ΔConcentration [Mm] | CRLB (%) | ΔConcentration [Mm] | CRLB (%) |
| 65-Gain | 0.03±0.03 | 2.53±0.05 | 0.03±0.02 | 46±2 | 0.05±0.02 | 6.13±0.09 |
| 50-Gain | -0.07±0.03 | 2.58±0.05 | 0.08±0.02 | 57±4 | 0.07±0.02 | 6.08±0.09 |
| 65-Loss | -0.02±0.04 | 2.62±0.05 | 0.04±0.02 | 45±2 | 0.06±0.02 | 6.07±0.07 |
| 50-Loss | 0.02±0.03 | 2.58±0.05 | 0.05±0.02 | 52±3 | 0.07±0.02 | 6.15±0.07 |

**Table S3. Mixed model 1 detailed results:** fixed effect coefficients (95% confidence intervals). LG- boolean loss/gain variable. GP- game probability, Boolean: 50% is 1, and 65% is 0.

| Variable | f value {Df1, Df2} | pValue |
|---|---|---|
| Intercept | 0.8 {1, 232.6} | 0.4 |
| ΔGlu | 46.7 {1, 380.4} | 3e-11 |
| LG | 3.2 {1, 292.0} | 0.07 |
| GP | 2.3 {1, 291.3} | 0.1 |
| LG*GP | 4.8 {1, 291.3} | 0.03 |

**Table S4. Mixed model 2 (Gain) detailed results**: fixed effect coefficients (95% confidence intervals). GP- game probability, Boolean: 50% is 1, and 65% is 0.

| Variable | f value {Df1, Df2} | pValue |
|---|---|---|
| Intercept | 769 {1, 47} | 9e-31 |
| ΔGABA | 5.8 {1, 47} | 0.03 |
| ΔGlu | 0.04 {1, 47} | 0.8 |

**Table S5. Mixed model 2 (Loss) detailed results**: fixed effect coefficients (95% confidence intervals). GP- game probability, Boolean: 50% is 1, and 65% is 0.

| Variable | f value {Df1, Df2} | pValue |
|---|---|---|
| Intercept | 650 {1, 47} | 3e-29 |
| ΔGABA | 0.3 {1 ,47} | 0.6 |
| ΔGlu | 0.007 {1, 47} | 0.5 |

**Table S6. Group-level activation maps clusters.** Gain- group level activation map for 65-Gain condition. Loss - group level activation map for 65-Loss condition. Learning covariate - group level activation map for Learning-GL map.

| Voxels | Region | Z-MAX | Z-MAX X (mm) | Z-MAX Y (mm) | Z-MAX Z (mm) |
|---|---|---|---|---|---|
| Gain-65 | | | | | |
| 224026 | occipital lobe | 10.3 | 45 | -71 | -4 |
| 16690 | Paracingulate gyrus | 7.07 | -2 | 13 | 49 |
| 5716 | Central Opercular Cortex | 5.98 | 47 | 6 | 2 |
| 1765 | Supramarginal Gyrus, anterior division | 5.34 | 68 | -27 | 26 |
| 1328 | Left Frontal Pole | 5.27 | -47 | 41 | 20 |
| 1262 | right thalamus | 5.35 | 15 | -12 | 5 |
| 912 | left hippocampus | 5.32 | -22 | -28 | -7 |
| 648 | Right Thalamus | 5.2 | 22 | -26 | -6 |
| 499 | Postcentral Gyrus | 6.23 | 60 | -17 | 49 |
| 480 | left Frontal Pole | 4.81 | -37 | 43 | 39 |
| Loss-65 | | | | | |
| 346684 | Occipital lobe | 10.4 | 44 | -72 | -5 |
| 13103 | right temporal lobe | 7.67 | 38 | 11 | 7 |
| 7151 | Left dlPFC | 5.53 | -36 | 50 | 29 |
| 4930 | right thalamus | 7.41 | 19 | -30 | -1 |
| 3467 | Right dlPFC | 5.58 | 32 | 45 | 30 |

**Table S7. Mixed model 3 (Gain) detailed results**: fixed effect coefficients (95% confidence intervals). GP- game probability, Boolean: 50% is 1, and 65% is 0.

| Variable | f value {Df1, Df2} | pValue |
|---|---|---|
| Intercept | 3.3 {1, 47} | 0.07 |
| ΔGABA | 0.03 {1, 47} | 0.8 |
| ΔGlu | 1.8 {1, 47} | 0.2 |
| Learning score | 9.1 {1, 47} | 0.004 |

**Table S8. Mixed model 3 (Loss) detailed results**: fixed effect coefficients (95% confidence intervals). GP- game probability, Boolean: 50% is 1, and 65% is 0.

| Variable | f value {Df1, Df2} | pValue |
|---|---|---|
| Intercept | 0.23 {1, 47} | 0.6 |
| ΔGABA | 1.7 {1, 47} | 0.2 |
| ΔGlu | 0.03 {1, 47} | 0.8 |
| Learning score | 2.2 {1, 47} | 0.1 |

**Table S9. Group-level learning correlation activation maps clusters.** Learning covariate - group level activation map for Learning-GL map.

| Voxels | Region | Z-MAX | Z-MAX X (mm) | Z-MAX Y (mm) | Z-MAX Z (mm) |
|---|---|---|---|---|---|
| Learning covariate: Gain>Loss | | | | | |
| 7302 | Left Postcentral Gyrus | 6.23 | -55 | -26 | 53 |
| 5389 | right occipital lobe | 5.26 | 15 | -59 | -12 |
| 1716 | left Lateral Occipital Cortex, inferior division | 6.26 | -37 | -88 | 0 |
| 1586 | right Lateral Occipital Cortex, superior division | 5.12 | 32 | -68 | 37 |
| 1363 | Left Thalamus | 4.57 | -13 | -18 | 3 |
| 1226 | left cingulate cortex | 4.96 | -6 | 5 | 45 |
| 1169 | left Precentral Gyrus | 6.26 | -32 | -7 | 72 |
| 1127 | left dlPFC | 5.16 | -27 | 55 | 27 |
| 985 | brain stem | 4.38 | 4 | -33 | -30 |
| 959 | left Lateral Occipital Cortex, superior division | 5.02 | -28 | -69 | 25 |
| 907 | Lingual Gyrus | 5.22 | -10 | -47 | 0 |
| 890 | right Intracalcarine Cortex | 5.29 | 8 | -80 | 10 |
| 696 | left occipital pole | 7.29 | -15 | -98 | -1 |
| 642 | left Lateral Occipital Cortex, inferior division | 4.55 | -48 | -65 | 0 |
| 570 | left cerebral cortex | 4.09 | -31 | -52 | -26 |
| 554 | right Thalamus | 4.19 | 15 | -28 | -3 |
| 535 | right temporal cortex/ anterior Supramarginal Gyrus | 6.11 | 67 | -19 | 42 |
| 509 | right Lateral Occipital Cortex, | 4.58 | 59 | -63 | 14 |
| 506 | right thalamus | 4.18 | 12 | -16 | 7 |
| 488 | cerebellum | 4.18 | 4 | -57 | -31 |

**Table S10. Mixed model 4 (Gain) detailed results**: fixed effect coefficients (95% confidence intervals). GP- game probability, Boolean: 50% is 1, and 65% is 0.

| Variable | f value {Df1, Df2} | pValue |
|---|---|---|
| Intercept | 14.5 {1, 60.7} | 0.0003 |
| ΔGABA | 5.1 {1, 77.4} | 0.03 |
| ΔGlu | 0.4 {1 75.5} | 0.5 |
| GP | 0.8 {1 35.7} | 0.4 |
| ΔGABA*GP | 0.6 {1 38.1} | 0.4 |
| ΔGlu*GP | 1.5 {1 38.4} | 0.2 |

**Table S11. Mixed model 4 (Loss) detailed results**: fixed effect coefficients (95% confidence intervals). GP- game probability, Boolean: 50% is 1, and 65% is 0.

| Variable | f value {Df1, Df2} | pValue |
|---|---|---|
| Intercept | 11.4 {1 91.5} | 0.001 |
| ΔGABA | 1.0 {1 92.6} | 0.3 |
| ΔGlu | 0.02 {1 92.9} | 0.9 |
| GP | 0.4 {1 45.7} | 0.5 |
| ΔGABA*GP | 1.3 {1 68.0} | 0.3 |
| ΔGlu*GP | 0.6 {1 62.2} | 0.4 |

Table S12. Connectivity Z value between dACC and other decision-making-related regions. Connectivity Z value between the dACC-functional MRS voxel and the listed brain regions for the fMRI gain group and the fMRI loss group. Bolded values represent significant values. HOSSA- Harvard-Oxford subcortical structure atlas. Amyg – Amygdala. Nuc.Acc- Nucleus Accumbens. Subcall. - Subcallosal cortex

| Mask location | Connectivity Z value (mean± SEM) (Gain group) | Connectivity Z value (mean± SEM) (Loss group) | Cohens d Effect Size (Gain group) | Cohens d Effect Size (Loss group) | Origin |
|---|---|---|---|---|---|
| Left Putamen | 0.334 ±0.043 | **1.072** | 0.296 ±0.037 | **1.084** | HOSSA |
| Right Putamen | 0.351 ±0.040 | **1.185** | 0.325 ±0.041 | **1.081** | HOSSA |
| Left Putamen | 0.378 ±0.045 | **1.155** | 0.335 ±0.037 | **1.222** | GL map |
| Right Putamen | 0.561 ±0.038 | **1.992** | 0.532 ±0.041 | **1.763** | GL map |
| Left Amyg. | 0.132 ±0.037 | 0.491 | 0.106 ±0.035 | 0.411 | HOSSA |
| Right Amyg. | 0.116 ±0.037 | 0.436 | 0.088 ±0.032 | 0.372 | HOSSA |
| Left insula | 0.698 ±0.042 | **2.248** | 0.686 ±0.034 | **2.735** | HOSSA |
| Right Insula | 0.736 ±0.038 | **2.620** | 0.742 ±0.038 | **2.656** | HOSSA |
| Left insula | 0.819 ±0.041 | **2.730** | 0.808 ±0.034 | **3.283** | GL map |
| Right Insula | 0.894 ±0.034 | **3.604** | 0.922 ±0.043 | **2.896** | GL map |
| Left Nuc.Acc | 0.072 ±0.028 | 0.358 | -0.030 ±0.025 | -0.160 | HOSSA |
| Right Nuc.Acc | 0.043 ±0.026 | 0.225 | -0.012 ±0.025 | -0.065 | HOSSA |
| Left caudate | -0.008 ±0.036 | -0.031 | -0.029 ±0.033 | -0.120 | GL map |
| Right Caudate | 0.008 ±0.039 | 0.027 | -0.006 ±0.035 | -0.021 | GL map |
| Left dlPFC | 0.340 ±0.034 | **1.347** | 0.336 ±0.042 | **1.093** | HOSSA |
| Right dlPFC | 0.434 ±0.049 | **1.215** | 0.479 ±0.047 | **1.387** | HOSSA |
| Left Thalamus | 0.199 ±0.042 | 0.645 | 0.209 ±0.031 | 0.925 | HOSSA |
| Right Thalamus | 0.261 ±0.040 | 0.891 | 0.209 ±0.034 | 0.847 | HOSSA |
| Left PCC | -0.026 ±0.040 | -0.089 | 0.069 ±0.040 | 0.232 | HOSSA |
| Right PCC | 0.085 ±0.036 | 0.320 | 0.148 ±0.045 | 0.451 | HOSSA |
| Left Subcall. | -0.073 ±0.037 | -0.269 | -0.091 ±0.028 | -0.453 | HOSSA |
| Right Subcall. | -0.096 ±0.037 | -0.360 | -0.107 ±0.028 | -0.514 | HOSSA |
| Left OFC | 0.007 ±0.039 | 0.023 | -0.004 ±0.035 | -0.016 | HOSSA |
| Right OFC | 0.073 ±0.045 | 0.219 | 0.081 ±0.041 | 0.271 | HOSSA |

**Table S13. A simulated likelihood ratio test to compare modeling using ΔGABA and ΔE/I balance.** A simulated likelihood ratio test compares the mixed models 1-4 containing ΔGABA and the same models with ΔE/I balance instead of ΔGABA. BIC- Bayesian Information Criterion. The comparison is made by the MATLAB command compare: compare (model_fit_GABA, model_fit_EI, 'nsim,' 1000). Number of simulation- 1000, alpha:0.05. AIC - Akaike Information Criterion.

| Mixed Model index | BIC {Df } | AIC | pValue |
|---|---|---|---|
| Model1-ΔGABA | -395.38 {7} | -423.16 | 1 |
| Model1-ΔE/I | 1617.2 {7} | 1589.4 | |
| Model2 Gain-ΔGABA | -20.834 {5} | -30.084 | 0.89 |
| Model2 Gain-ΔE/I | -19.031 {5} | -28.282 | |
| Model2 Loss-ΔGABA | -9.4495 {5} | -18.7 | 0.72 |
| Model2 Loss-ΔE/I | -9.1588 {5} | -18.41 | |
| Model3 Gain-ΔGABA | 152.82 [1] | 141.71 | 0.33 |
| Model3 Gain-ΔE/I | 152.72 {6} | 141.62 | |
| Model3 Loss-ΔGABA | 140.39 {6} | 129.29 | 0.80 |
| Model3 Loss-ΔE/I | 141.44 {6} | 130.33 | |
| Model4 Gain-ΔGABA | 271.9 {8} | 251.73 | 0.55 |
| Model4 Gain-ΔE/I | 272.37 {8} | 252.19 | |
| Model4 Loss-ΔGABA | 297.95 {8} | 277.69 | 0.72 |
| Model4 Loss-ΔE/I | 298.66 {8} | 278.4 | |

**Table S14. $T_2$ Values.** $T_2$ literature values were used for transverse relaxation correction of the quantified LCM results.

| Metabolite | water | NAA/NAAG/tNAA | Cr/pCr/tCr | Cho/GPC/pCh/tCho | mI | Glu/Gln/Glx | Tau | GSH | GABA |
|---|---|---|---|---|---|---|---|---|---|
| $T_2$ [ms] | 47 | 132 | 95 | 152 | 95 | 93 | 93 | 61 | 87 |
| Brain area | Occipital lobe | | | | | | | | |
| Voxel Size [ml] | 19.7 | | | | | | | | 27 |
| Tissue Fraction | GM/WM/CSF- 0.51/0.44/0.05 | | | | | | | | |
| Reference | (Marjańska et al., 2012)(1) | | | | | | | | (Andreychenko et al., 2013)(2) |

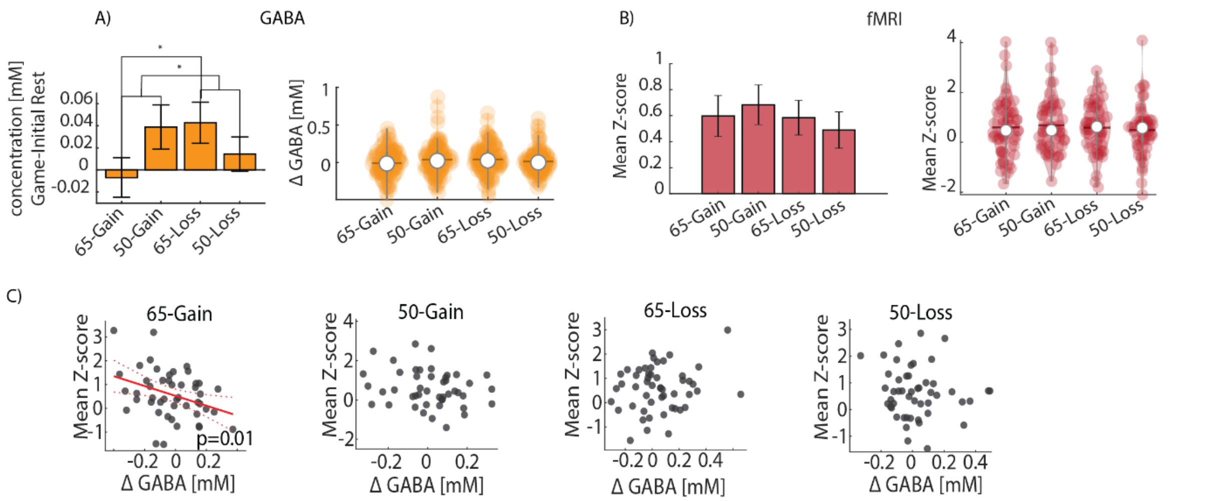


**Figure S1. ΔGABA and BOLD measurements and correlation under all game conditions. A)** ΔGABA concentration and distribution during all game conditions. **B)** fMRI mean Z score and distribution during all game conditions. **C)** Correlation between BOLD activities in mean Z score from the dACC cluster in the ROI-GL map and ΔGABA during all games.

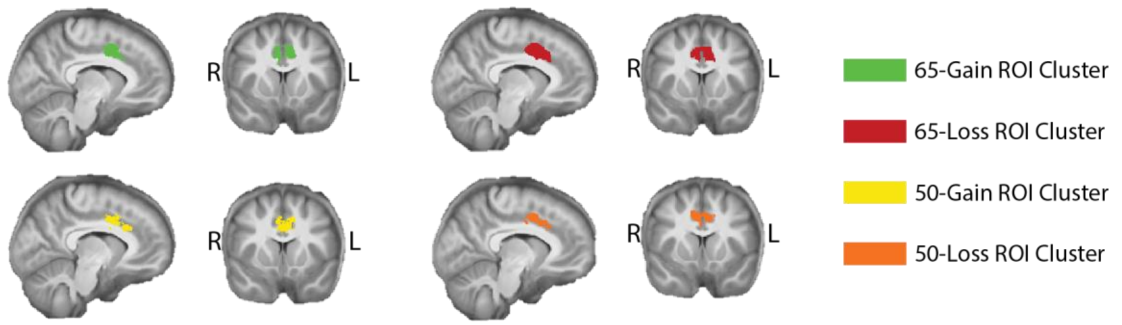


**Figure S2. Group-level analysis results using the spectroscopic voxel as a region of interest (ROI).** Contrast: decision phase (coding of value difference). Images are z-scored, with 3.1 thresholds.

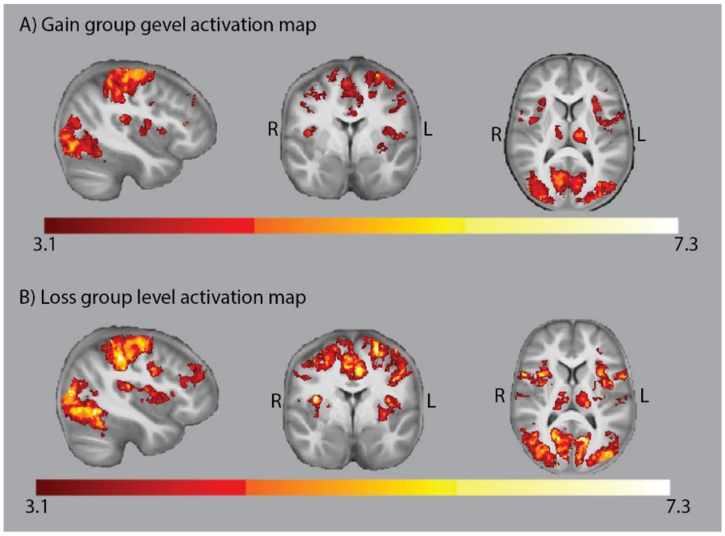


**Figure S3. Group level analysis results of 65-35 games.** Contrast: decision phase (coding of value difference). Images are z-scored, with 3.1 threshold. A) Gain group. B) Loss

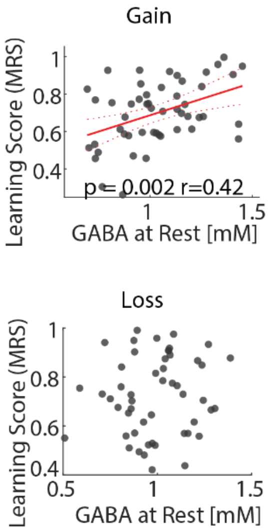


**Figure S4. Learning correlation with rest compared to 65-gain levels of GABA.** correlation between GABA concentration at rest and learning scores during the gain and loss 65-35 games.
